# The molecular determinants of *R*-roscovitine block of hERG channels

**DOI:** 10.1101/644336

**Authors:** Bryan Cernuda, Christopher Fernandes, Salma Allam, Matthew Orzillo, Gabrielle Suppa, Zuleen Chia Chang, Demosthenes Athanasopoulos, Zafir Buraei

## Abstract

<u>H</u>uman <u>e</u>ther-à-go-go-<u>r</u>elated <u>g</u>ene (Kv11.1, or hERG) is a potassium channel that conducts the delayed rectifier potassium current (I_Kr_) during the repolarization phase of cardiac action potentials. hERG channels have a larger pore than other K^+^channels and can trap many unintended drugs, often resulting in acquired LQTS (aLQTS). *R*-roscovitine, a cyclin-dependent kinase (CDK) inhibitor that also inhibits L-type calcium channels, inhibits open hERG channels but does not become trapped in the pore. Two-electrode voltage clamp recordings from *Xenopus* oocytes expressing wild-type (WT) or mutant (T623A, S624A, Y652A, F656A) hERG channels demonstrated that, compared to WT hERG, T623A, Y652A, and F656A inhibition by 200 μM *R*-roscovitine was ~ 48 %, 29 %, and 73 % weaker, respectively. In contrast, S624A hERG was inhibited more potently than WT hERG, with an ~ 34 % stronger inhibition. These findings were further supported by the IC_50_ values, which were increased for T623A, Y652A and F656A (by ~5.5, 2.75, and 42 fold respectively) and reduced 1.3 fold for the S624A mutant. Our data suggest that while T623, Y652, and F656 are critical for *R*-roscovitine-mediated inhibition, S624 may not be. This relatively unique feature, coupled with *R*-roscovitine’s tolerance in clinical trials, could guide future drug screens. We discuss our findings and how they lend support for the recent Comprehensive *In Vitro* Proarrhythmia Assay (CiPA) guidelines on the re-evaluation of potentially useful drugs that had failed testing due to unintended interactions with hERG.

## Introduction

Human ether-à-go-go-related gene, or hERG [Kv11.1], is a voltage-gated potassium channel critical for nerve and cardiac function [1,2]. In the heart, hERG channels initially open during the depolarization phase of the cardiac action potential (cAP) but immediately inactivate. Upon cAP repolarization, hERG channels de-inactivate and reopen, which allows the ensuing large K^+^ efflux to speed cAP repolarization [1], limit cardiac excitability, and maintain normal QT intervals [3]. Consequently, mutations in hERG are one of the leading causes of congenital long QT syndrome (cLQTS), with a neonatal incidence rate of up to 1 in 2,500 [4]; abnormal cardiac phenotypes are usually triggered during exercise, arousal, or rest [5].

hERG channels are tetrameric proteins, with each monomer consisting of six transmembrane alpha helices (S1-S6) and cytoplasmic amino and carboxy termini [6]. Similar to other voltage-gated channels, S1-S4 is considered the primary voltage-sensing region, with S4 containing positively-charged residues that move slowly outward to induce the characteristically slow activation kinetics of hERG [7,8]. The four S5 and S6 helices and their intervening sections, including the P-loops and P-helices, form the pore and selectivity filter of the channel [9,10]. The pore is thought to have a region between the pore helix and the S6 segments that may provide pockets for drug binding [11]. Together with a strong negative electrostatic potential, the hERG pore becomes highly susceptible to unwanted drug interactions, which can result in current inhibition and acquired long QT syndrome (aLQTS) [12]. aLQTS greatly predisposes individuals to lethal cardiac arrhythmias [13], and is by far more common than cLQTS. It is estimated that 1 out of every 5 intensive care unit (ICU) patients experiences a form of cardiac arrhythmia that is highly influenced by QT-prolonging medications [14–16]. Because of the dangerous side effects, pharmaceuticals have been routinely screened for hERG interaction [17,18], which has unfortunately led to the loss of promising drugs that may not have posed a clinical proarrhythmic risk [19]. Interestingly, many clinically-safe drugs turned out to inhibit hERG, which prompted CiPA guidelines recommending the re-evaluation of previously discarded, yet promising drugs; the features of such drugs are under investigation [19].

*R*-roscovitine is a cyclin-dependent kinase (CDK) inhibitor that has been used in clinical trials for cancer and for Cushing’s disease [20,21]. We previously found that *R*-roscovitine also blocks voltage-gated K^+^channels by binding to the open state [22]. This interaction may partially contribute to *R*-roscovitine’s anticancerogenic effects since voltage-gated K^+^channels, including hERG channels, are overexpressed in many cancer cell lines and are linked to tumorigenesis [23,24]. *R*-roscovitine inhibits hERG channels in a unique fashion [25]: 1) it blocks open hERG channels and does not become trapped in the pore during channel inactivation or closing, 2) it has low or no preference to closed and inactivated states, and 3) it does not exhibit use-dependence, suggesting very rapid block and unblock kinetics [25]. These distinct characteristics of *R*-roscovitine, as well as the absence of arrhythmic side effects during clinical trials [20], alluded to a potentially unique binding mechanism for *R*-roscovitine compared to other hERG inhibitors. Thus, we set out to investigate the hERG binding determinants for *R*-roscovitine.

Using electrophysiological recordings on mutant hERG channels, we found that two hERG residues (S624 and Y652) that, otherwise concordantly contribute to drug interactions [26], have opposing effects on *R*-roscovitine mediated inhibition. Specifically, hERG Y652A resisted *R*-roscovitine inhibition while hERG S624A currents were strongly inhibited, significantly more so than WT currents. We also characterized two other residues frequently involved in binding to hERG blockers: a residue adjacent to the selectivity filter (T623) and another S6 helix residue (F656). The mutation of either residue significantly weakened *R*-roscovitine block, suggesting the importance of both residues. These molecular determinants highlight a complexity that may guide future work on open state block specificity. In addition, our results exemplify the need to use CiPA guidelines when assessing clinically promising drugs [19].

## Methods

### RNA Preparation and Oocyte Injection

WT and mutant (T623A, S624A, Y652A, and F656A) hERG clones, which were subcloned into a pSP64 vector, were ordered from Addgene (catalog #53051 for WT, #53055 for T623A, #53056 for S624A, #53053 for Y652A, and #53054 for F656A). The vectors were linearized with EcoRI and *in vitro* transcribed using the mMESSAGE mMACHINE SP6 Transcription Kit (Ambion). 1.0 μg of the resulting cRNA was injected (using Nanoject II from Drummond Scientific, with Drummond Scientific glass capillaries) into defolliculated *Xenopus laevis* oocytes that were generously provided by Dr. Jian Yang and Yong Yu (Columbia University), as well as purchased from Xenopus 1. Oocytes (stages V-VI) were prepared as previously described [27]. Briefly, 2-3 mm size strips from *Xenopus laevis* ovaries were treated with a solution containing 0.3-0.5 mg/ml collagenase A (Boehringer Mannheim), 82.5 mM NaCl, 2.5 mM KCl, 1 mM MgCl_2_, and 5 mM HEPES (pH 7.6) for ~ 1.5 hours on a shaking incubator set to 175 rpm. A 15-minute rinse on a shaking incubator set to 75 rpm was repeated twice with ND96 solution that contained 96 mM NaCl, 2.5 mM KCl, 1 mM MgCl_2_, 5 mM HEPES, 1.8 mM CaCl_2_, 100 units/mL penicillin, 100 μg/mL streptomycin (pH 7.6), and 5 mM sodium pyruvate (used only with eggs ordered from Xenopus 1). Following the selection of defoliculated oocytes, hERG cRNAs were injected into the oocytes and recordings were performed 2-7 days later. All procedures were in accordance with regulations set by the institutional animal care and use committee.

### Solutions and drug administration

Electrodes (Clark capillary glass, number 30-0038, Harvard Apparatus) were pulled on a Narishige PC-10 puller, and used with a resistance of 1-5 MΩ. The intracellular electrode solution contained 3M KCl, while the low potassium extracellular (recording) solution contained [in mM]: 5 KCl, 100 NaCl, 1.5 CaCl_2_, 2 MgCl_2_, 10 HEPES. To elicit large inward tail currents for hERG mutants with reduced permeation, a high potassium extracellular solution was used, which contained [in mM]: 96 KCl, 2 NaCl, 1.8 CaCl_2_, 1 MgCl_2_, 5 HEPES. The pH of the external solutions were adjusted to 7.4 using either 1M NaOH (for low K^+^) or 1M KOH (for high K^+^), and the osmolarities were 200 mOsm. *R*-roscovitine was dissolved in DMSO to make a 100 mM stock solution that was frozen at −80°C. Drug-free external solutions (control and wash) had the same DMSO concentrations as the *R*-roscovitine-containing solutions. Working solutions were prepared on the day of the recordings.

### Two-Electrode Voltage Clamp (TEVC) Recordings

Currents were amplified using the OCT clamp TEVC instrument and data was acquired and analyzed using Clampex and Clampfit, respectively (Axon Instruments). Currents were digitized using a digidata 1440A board at 10 kHz. Leak subtraction was not applied during the recordings. Before recordings began, a pulse protocol was applied to allow hERG runup or rundown to subside. The pulse protocol consisted of a −80 mV holding potential, a 1-second depolarization to +40 mV, and a 1-second repolarization to either −50 mV (for low K^+^ external solutions) or −120 mV (for high K^+^ external solutions). The protocol was applied for ~ 3.5 minutes. Following this period, other voltage protocols [described below] were applied. All traces were baseline-adjusted to remove the minuscule current that deviated from 0 μA at the −80 mV holding potential.

### Time-Course Protocol and IC_50_ Calculations

Cells were held at −80 mV. For outward-current hERG constructs, a 1-second depolarizing step from −80 mV to +40 mV was followed by a 1-second repolarization to −50 mV. This pulse was applied repetitively over a span of ~ 33 minutes. A similar protocol was applied to inward-current hERG constructs, but the 1-second repolarization voltage was instead set to −120 mV. Tail currents were measured to generate a time-course of inhibition for each cell before and during application of various *R*-roscovitine concentrations: 10, 30, 100, 300, and 1000 μM. Fractional block of tail current was calculated for each *R*-roscovitine concentration as: (*I*_control_−*I*_Rosc_)/*I*_control_, with *I*_control_ being an average of the tail current before a particular *R*-roscovitine concentration and the maximum drug-free current elicited from the cell [i.e. (*I*_before_ + *I*_maxbefore_)/2]. The fractional block was then plotted against *R*-roscovitine concentrations and fitted with the following Hill equation to obtain IC_50_ values: Y = Bottom+(Top-Bottom)/(1+10^((LogIC_50_−X)*HillSlope)), where X is the log of the concentration, Top and Bottom (constrained to 1 and 0, respectively) are the curve plateaus, LogIC_50_ is the center of curve, and HillSlope is the slope factor.

### Step Depolarization and Step Repolarization Protocols

Cells were held at −80 mV and 2-second voltage steps were applied in 10 mV increments from −50 mV to +60 mV. The resulting step currents were measured as current means from the final 50 ms of the depolarization, which were then normalized and plotted against step voltages to generate step I-V curves. Following the depolarizing steps, cells were repolarized to −50 mV for 1.5 seconds, eliciting tail currents. Peak tail currents were normalized and plotted against step voltages for tail I-V curves, which were fitted with the following Boltzmann sigmoidal equation to obtain activation curves: Y = Bottom+(Top-Bottom)/(1+exp((V_50_−X)/Slope)), where Top and Bottom are the curve plateaus (constrained to 1 and 0, respectively), V_50_ is the step voltage that elicits 50% channel activation, and Slope is the steepness of the curve. Percent inhibition of step and tail currents with 200 μM *R*-roscovitine was calculated as: (*I*_control_−*I*_Rosc,200μM_)/*I*_control_, with *I*_control_ being the current size before 200 μM *R*-roscovitine application.

A step repolarization protocol was used for inward-current hERG constructs. Following 1-second depolarization steps to +60 mV, 2-second repolarization voltages were applied in 20 mV decrements from +20 mV to −120 mV. The resulting peak tail currents were then normalized and plotted against tail voltages to acquire tail I-V curves, which were fitted with a second-order polynomial equation (quadratic): Y = B0+B1*X+B2*X^2^. To obtain reversal potentials (*E*_rev_), the equation was solved for Y = 0. Similar to step and tail current inhibitions [during step depolarization protocols] with 200 μM *R*-roscovitine, percent inhibition of tail currents with 500 μM *R*-roscovitine was calculated as: (*I*_control_−*I*_Rosc,500μM_)/*I*_control_, with *I*_control_ being the current size before 500 μM *R*-roscovitine application.

### Data Analysis and Statistics

Calmpex was used for data acquisition and Clampfit and Microsoft Excel were used for initial data analyses. GraphPad Prism version 8.0.2 for Macintosh (GraphPad Software, La Jolla California USA) was used to make I-V, time-course, and percent inhibition graphs; it also calculated the nonlinear fits of the tail I-V and fractional block graphs (i.e. Boltzmann, four parameter logistic, and quadratic). Data are represented as mean ± standard error. All statistical analyses were performed using GraphPad Prism. Ordinary one-way ANOVA tests and Kruskal-Wallis tests were followed by post-hoc analyses: Dunnett’s multiple comparison test [for one-way ANOVAs] and Dunn’s test [for Kruskal-Wallis tests], with multiple comparisons reported as multiplicity adjusted p values. Statistical significance was designated when p < 0.05 and all p-values were subsequently reported.

## Results

### *R*-roscovitine-mediated inhibition of WT hERG in Xenopus oocytes

To determine the appropriate concentration of *R*-roscovitine for comparing different hERG constructs, we wanted to establish the effectiveness of *R*-roscovitine inhibition on wild-type (WT) hERG channels in *Xenopus* oocytes. The half-maximal inhibitory concentration (IC_50_) of tail currents for *R*-roscovitine with WT hERG was determined previously in HEK-293 cells (27 μM), but *Xenopus* oocytes are known to require higher drug doses [25]. Therefore, we injected cRNA for WT hERG into *Xenopus* oocytes and recorded potassium currents 2-7 days later (see methods). The currents were elicited using a 2-second pulse protocol consisting of a 1-second depolarization to +40 mV followed by a 1-second repolarization to −50 mV, during which tail currents were measured (Fig 1B). This pulse protocol was repeated either every 5 seconds for *R*-roscovitine concentrations applied in a randomized order (100, 300, 30, and 10 μM) or every 12 seconds for ascending *R*-roscovitine concentrations (10, 30, 100, 300, and 1000 μM), with the various drug concentrations being repeatedly applied and washed off (Fig 1C). Fitting the fractional block of hERG tail currents with a Hill equation resulted in an IC_50_ for WT of 196 ± 12 μM (Fig 1D). Thus, we used 200 μM *R*-roscovitine for studying inhibition levels of WT and mutant hERG channels.

**Fig 1.**
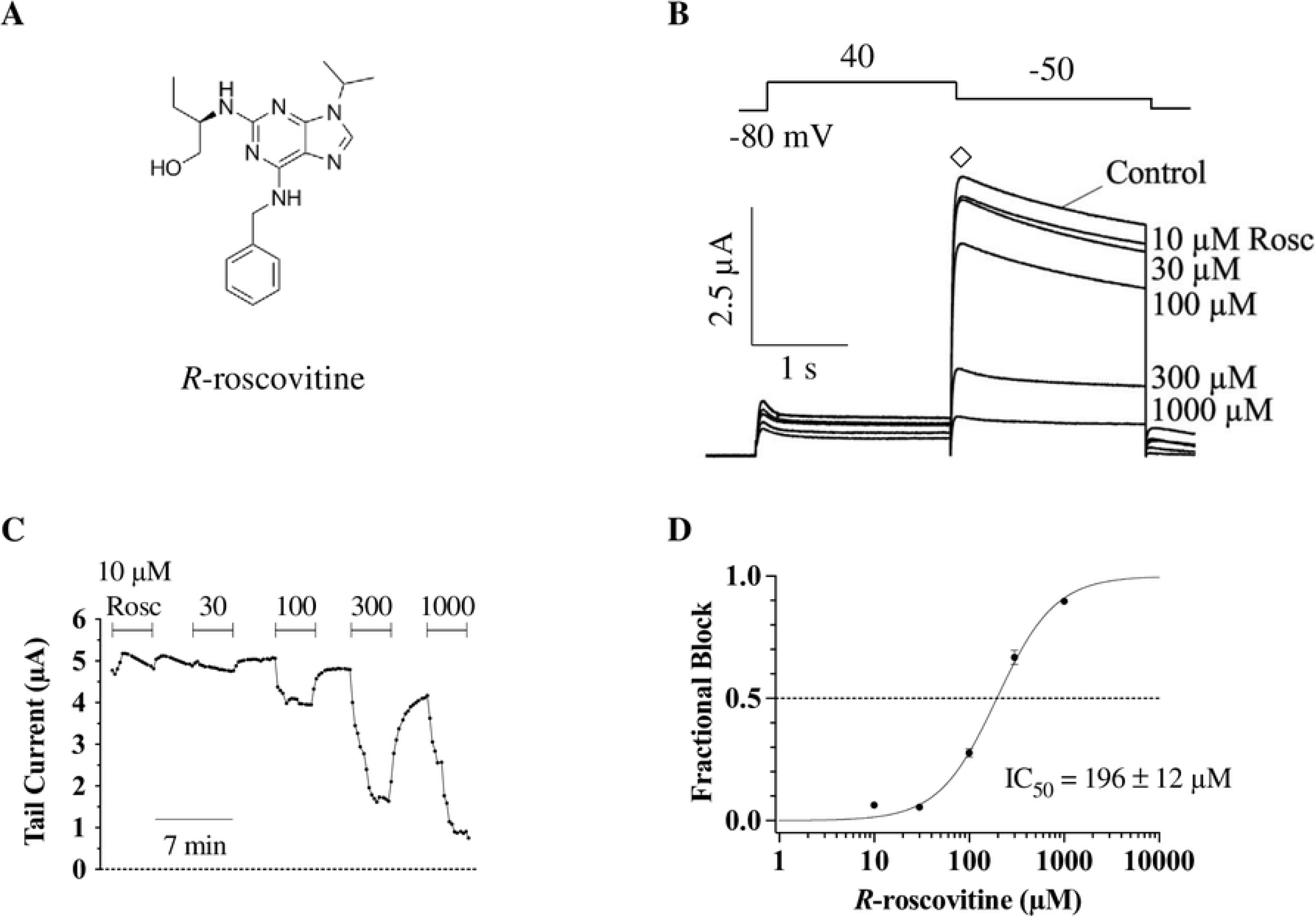
IC_50_ value for *R*-roscovitine binding to WT hERG channels. A) Skeletal formula of *R*-roscovitine, 2-(1-ethyl-2-hydroxyethylamino)-6-benzylamino-9-isopropylpurine. B) Pulse protocol (top) along with representative traces from a WT cell (bottom) exposed to the indicated *R*-roscovitine concentrations (10, 30, 100, 300, and 1000 μM). Channels were activated with a 1-second step to +40 mV from −80 mV, and tail currents (♢) were elicited by a 1-second repolarization to −50 mV. The single 2-second episodic pulse was repeated for 3.44 minutes as the various concentrations were washed in and out of the gravity-fed perfusion system. C) Time-course of tail current inhibition (measured from ♢, in B) from a representative cell expressing WT hERG in the presence of the indicated *R*-roscovitine concentrations. D) Fractional block data were fitted with a Hill equation (see methods) to obtain the IC_50_ for WT hERG (196 ± 12 μM, n = 11). Error bars here and elsewhere represent standard errors.

We next compared currents in the presence and absence of the drug during a step depolarization protocol. Specifically, step currents were elicited using a series of 2-second depolarizations ranging from −50 mV to +60 mV (in 10 mV increments), and the subsequent tail currents were recorded during a 1.5-second repolarization step to −50 mV (Fig 2A, n = 14). Typical n-shaped current-voltage (I-V) curves for WT hERG were obtained when step currents were measured at the end of the depolarizing voltages (Fig 2B); the current decline with depolarizing potentials was due to the fast inactivation at these voltages [1,28]. As expected based on a previous study, where *R*-roscovitine inhibition occurred only with open hERG channels [25], 200 μM *R*-roscovitine inhibition was weak at both lower and higher voltages (e.g. 23.4 ± 4.5 % at −50 mV and 8.2 ± 5.4 % at +50 mV); the diminished inhibition was due to low open probability at low voltages and a strong inactivation at high voltages. Inhibition was strongest at intermediate voltages [where inactivation was weak and open probability high], reaching a maximum of 43.3 ± 3.0 % at 0 mV (p = 0.0052 for 0 vs. −50 mV, p < 0.0001 for 0 vs. +50 mV, n = 14, one-way ANOVA; Fig 2C).

**Fig 2.**
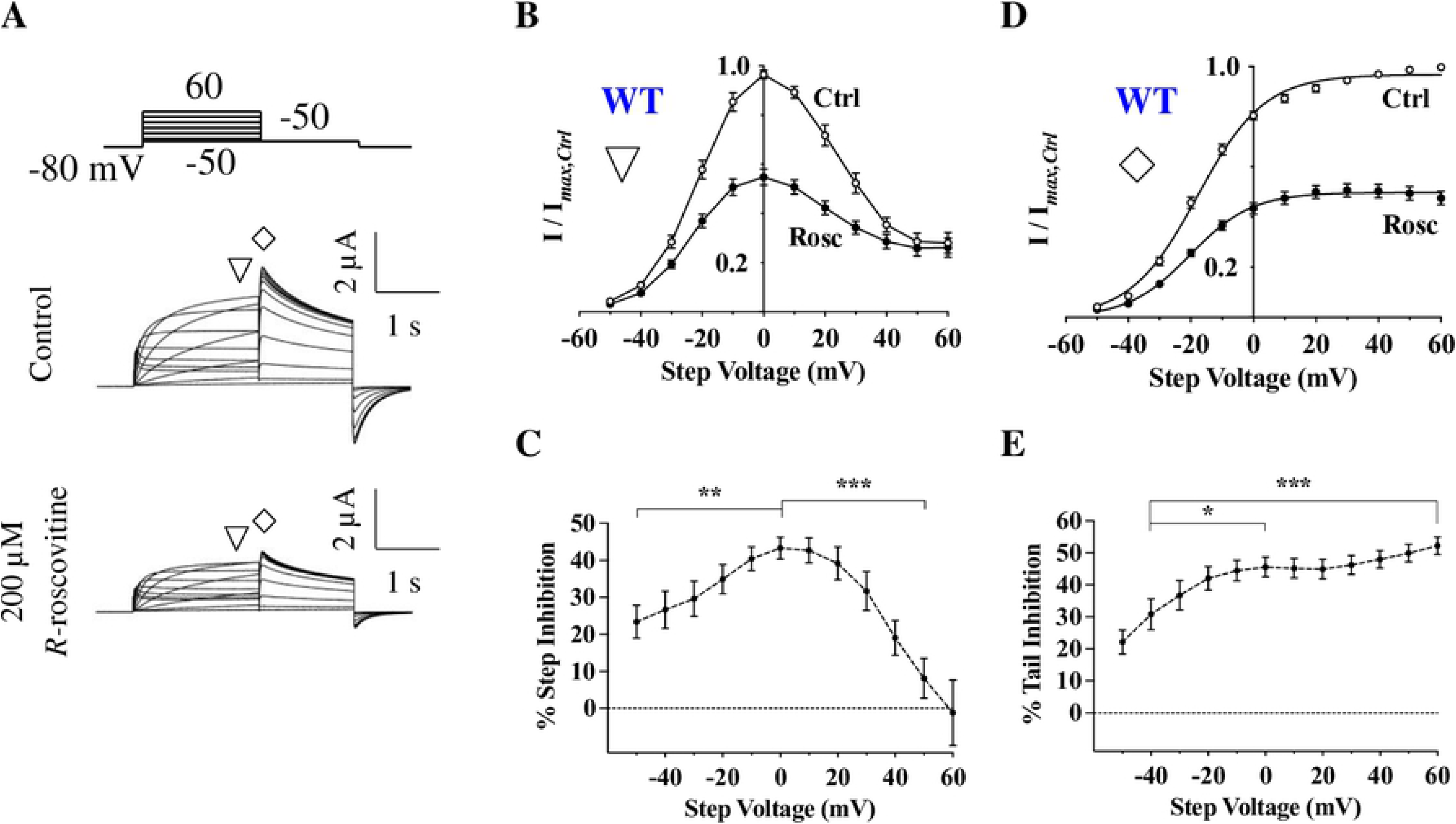
Characteristics of *R*-roscovitine inhibition of hERG channels. A) Voltage protocol and current traces from a representative cell expressing hERG channels before (Control) and during 200 μM *R*-roscovitine application. From a holding potential of −80 mV, outward currents were elicited at two phases: depolarization (from −50 mV to +60 mV) and repolarization (to −50 mV). Mean step currents were measured at the end of the 2-second step (∇) and tail currents were measured from their peak (♢). B) Normalized I-V curves for Control and 200 μM *R*-roscovitine from mean step currents (∇, n = 14). Inhibition was strongest between −30 mV and +30 mV, a range where the open state is more prevalent than either the closed or inactivated state. C) Average percent step current inhibition by 200 μM *R*-roscovitine at various step voltages. Inhibition was weakest at voltages where channels are closed (−50 mV) or inactivated (+50 mV; p = 0.0052 for 0 vs. −50 mV, p < 0.0001 for 0 vs. +50 mV, n = 14, one-way ANOVA). D) Tail I-V curves for Control and 200 μM *R*-roscovitine, with normalized peak tail currents (♢, n = 14). Smooth lines are Boltzmann fits to the activation curves, which produced V_0.5_ (Ctrl: −16.5 ± 1.1 mV, Rosc: −19.4 ± 1.1 mV; p = 0.0105 for Ctrl vs. Rosc, n = 14, paired *t*-test) and slopes (Ctrl: 11.7 ± 0.5, Rosc: 10.6 ± 0.5) of activation. E) Average percent tail current inhibition by 200 μM *R*-roscovitine at various step voltages. The inhibition of tail current significantly increased with rising levels of depolarized potentials (p = 0.0134 for −40 vs. 0 mV, p = 0.0004 for −40 vs. +60 mV, n = 14, one-way ANOVA). Error bars represent standard errors; some error bars are smaller than the symbols. * = *P* < 0.05, ** = *P* < 0.01, and *** = *P* < 0.001.

Since hERG channels inactivate very rapidly during step depolarizations, hERG channel block is normally measured from tail currents [29,30]. Following the depolarization step, repolarization to −50 mV allowed hERG channels to deinactivate rapidly and transition to the open state, causing a large efflux of K^+^ ions (Fig 2A); these tail currents reflect the proportion of channels activated at the preceding depolarized potentials [28,31]. Plotting normalized peak tail currents against step voltages resulted in typical sigmoidal activation curves (Fig 2D, n = 14), with a threshold for current activation of approximately −40 mV and a near-maximum channel activation at +40 mV. Activation curves were fitted with a Boltzmann equation, yielding a V_0.5_ of activation of −16.5 ± 1.1 mV. Application of 200 μM *R*-roscovitine shifted the V_0.5_ of activation by ~ −3 mV (V_0.5,*Rosc*_= −19.4 ± 1.1 mV, p = 0.0105 for Ctrl vs. Rosc, n = 14, paired *t*-test). In addition, *R*-roscovitine inhibition of tail current significantly increased with depolarization from 30.9 ± 4.9 % at −40 mV to 45.6 ± 3.1 % at 0 mV, and reached its maximum of 52.3 ± 2.7 % at +60 mV (p = 0.0134 for −40 vs. 0 mV, p = 0.0004 for −40 vs. +60 mV, n = 14, one-way ANOVA; Fig 2E). Thus, *R*-roscovitine inhibition of tail current increased with channel activation.

### Comparison of WT and mutant hERG channels in the absence of *R*-roscovitine

Next, we examined currents from hERG mutants to establish their baseline properties before applying *R*-roscovitine. hERG block is commonly associated with residues located near the selectivity filter (e.g. S624) and on the S6 helices (e.g. Y652) [32,33]. Therefore, single-mutant channels S624A and Y652A were expressed in *Xenopus* oocytes and subjected to a step depolarization protocol (Fig 3A, top). Step I-V curves for WT, S624A, and Y652A showed maximum step currents at 0 mV, −10 mV, and +10 mV, respectively (Fig 3A, bottom). At +60 mV, S624A had more than twice the relative amount of current compared to WT channels, which indicated possible incomplete inactivation; this mutant channel was previously characterized to have impaired inactivation [34]. Boltzmann fits to the activation curves indicated no significant difference in half-maximal activation between S624A (−17.7 ± 3.2 mV), Y652A (−16.5 ± 1.0 mV), and WT (−16.5 ± 1.1 mV) (p > 0.05 for WT vs. S624A and WT vs. Y652A, n ≥ 9 for all groups, one-way ANOVA; Fig 3B).

**Fig 3.**
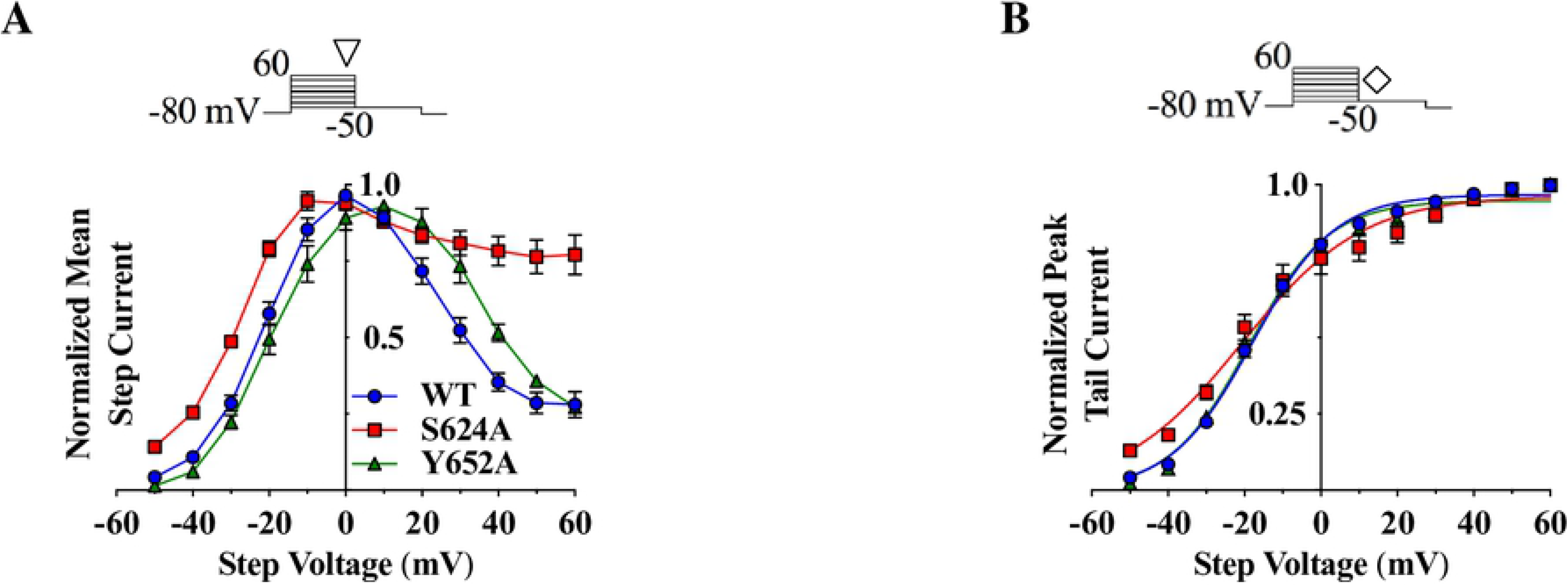
Comparison of unaffected step and tail currents from WT hERG, S624A, and Y652A hERG mutants. A) Step I-V curves for WT (ⵔ, blue), S624A (☐, red), and Y652A (Δ, green) from a step depolarization protocol (top; currents measured at ∇). Rectification of S624A currents was reduced in comparison to WT current, with more than twice the amount of normalized current remaining for S624A (0.77 ± 0.06) compared to WT (0.28 ± 0.04) at +60 mV. B) WT, S624A, and Y652A tail I-V curves from a step depolarization protocol (top; currents measured at ♢). Boltzmann fits generated the following V_0.5_ and slopes of activation: WT (−16.5 ± 1.1 mV, 11.7 ± 0.5), S624A (−17.7 ± 3.2 mV, 16.3 ± 1.6), and Y652A (−16.5 ± 1.0 mV, 12.5 ± 0.8). S624A and Y652A showed no significant differences in V_0.5_ of activation when compared to WT (p > 0.8 for both comparisons, n_WT_= 14, n_S624A_= 9, n_Y652A_= 9, one-way ANOVAs).

### The S624A mutation enhances *R*-roscovitine block

To determine how changes to the hERG pore residues alter *R*-roscovitine inhibition, WT and mutant channel currents were compared in the presence and absence of 200 μM *R*-roscovitine. In general, mutating key residues involved in drug-binding has been shown to weaken or abolish drug inhibition [35]. Applying the step depolarization protocol to S624A hERG channels resulted in currents with typical n-shaped step I-V curves and inhibition occurred at intermediate voltages (i.e. between −30 mV and +30 mV; Fig 4B). Surprisingly, mutating the S624 residue did not attenuate 200 μM *R*-roscovitine current inhibition: S624A and WT step current inhibition were almost identical over a range of step voltages (p > 0.05 for WT vs. S624A between +10 mV and +60 mV, n ≥ 9, one-way ANOVA; Fig 4C). In fact, at a few intermediate voltages, the inhibition of S624A was almost 25% larger than WT inhibition (p < 0.05 for WT vs. S624A between −20 mV and 0 mV, n ≥ 9, one-way ANOVA).

**Fig 4.**
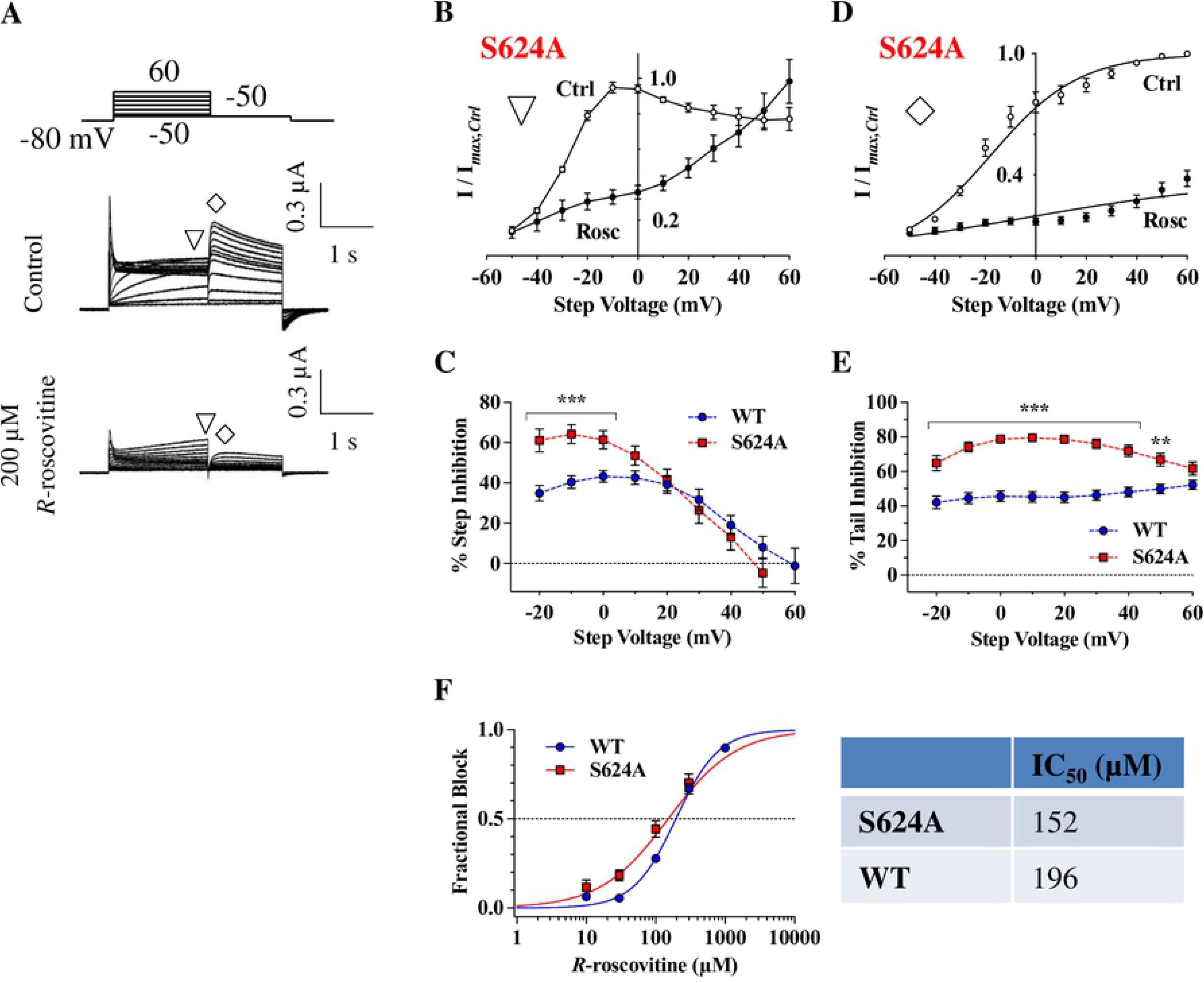
S624A hERG channels have an increased sensitivity to *R*-roscovitine. A) Voltage protocol and representative current traces of a cell before and during 200 μM *R*-roscovitine application. Note that tail currents were greatly diminished upon *R*-roscovitine application. B) S624A step I-V curves before and during 200 μM *R*-roscovitine application (currents measured at ∇). The curves had a similar bell shape to WT step I-V curves, with inhibition occurring between −30 mV and +30 mV. C) Percent step inhibition of WT and S624A at varying step voltages. S624A was inhibited more potently than WT at intermediate voltages, where open probability is high (p < 0.05 for WT vs. S624A between −20 mV and 0 mV, n_WT_= 14, n_S624A_= 9, one-way ANOVA). D) S624A tail I-V curves measured at ♢. Smooth lines are Boltzmann fits that generated V_0.5_ (Ctrl: −17.7 ± 3.2 mV, Rosc: −3.3 ± 6.1 mV) and slopes (Ctrl: 16.3 ± 1.6, Rosc: 44.5 ± 3.3) of activation. No significant shifts in V_0.5_ of activation occurred with drug application (p = 0.09, n = 9, paired *t*-test). E) Percent tail inhibition of WT and S624A. When compared to WT over a range of voltages, levels of S624A tail current inhibition were much larger (p < 0.05 for WT vs. S624A between −20 mV and +50 mV, n_WT_= 14, n_S624A_= 9, one-way ANOVA). F) Concentration-response relationship for S624A. Although the S624A IC_50_ (152 ± 22 μM) was lower than the WT IC_50_ (196 ± 12 μM), there was no statistical difference (p = 0.8323 for WT vs. S624A, n_WT_= 11, n_S624A_= 9, one-way ANOVA).

S624A tail I-V curves had typical sigmoidal curves and a V_0.5_ of activation of −17.7 ± 3.2 mV before *R*-roscovitine application (Fig 4D). Although there seemed to be a shift during *R*-roscovitine block (S624A V_0.5,*Rosc*_= −3.3 ± 6.1 mV), this shift was not statistically significant (p = 0.09, n = 9, paired *t*-test). Similar to the enhanced step inhibition at intermediate step voltages, S624A tail current inhibition was significantly stronger than WT channel inhibition over a range of depolarized voltages (p < 0.05 for WT vs. S624A between −20 mV and +50 mV, n ≥ 9, one-way ANOVA; Fig 4E). At +50 mV, S624A tail inhibition by 200 μM *R*-roscovitine reached 66.8 ± 3.6%, which was larger than WT inhibition (49.9 ± 2.8%) at the same step voltage (p = 0.0012, n ≥ 9, one-way ANOVA). When we measured dose-response curves, S624A IC_50_ (152 ± 22 μM) was smaller than WT IC_50_ (196 ± 12 μM), but this difference did not reach statistical signifiance (p > 0.05 for WT vs. S624A, n ≥ 9, one-way ANOVA; Fig 4F). The persistence of inhibition suggested that S624 may not be required for *R*-roscovitine inhibition. It remained unclear, however, as to why inhibition levels increased at some voltages in S624A hERG.

### The Y652A mutation weakens *R*-roscovitine inhibition of hERG channels

We next tested the importance of the Y652 residue, which is a key binding target for several hERG inhibitors [36]. Using the same step depolarization as before (Figs. 2, 3, 4), step and tail currents were elicited in the presence or absence of 200 μM *R*-roscovitine (Fig 5A). Unlike S624A, Y652A hERG exhibited a reduced inhibition of step current compared to WT hERG (Fig 5B, C; p < 0.05 for WT vs. Y652A between −20 mV and +10 mV, n ≥ 9, one-way ANOVA).

**Fig 5.**
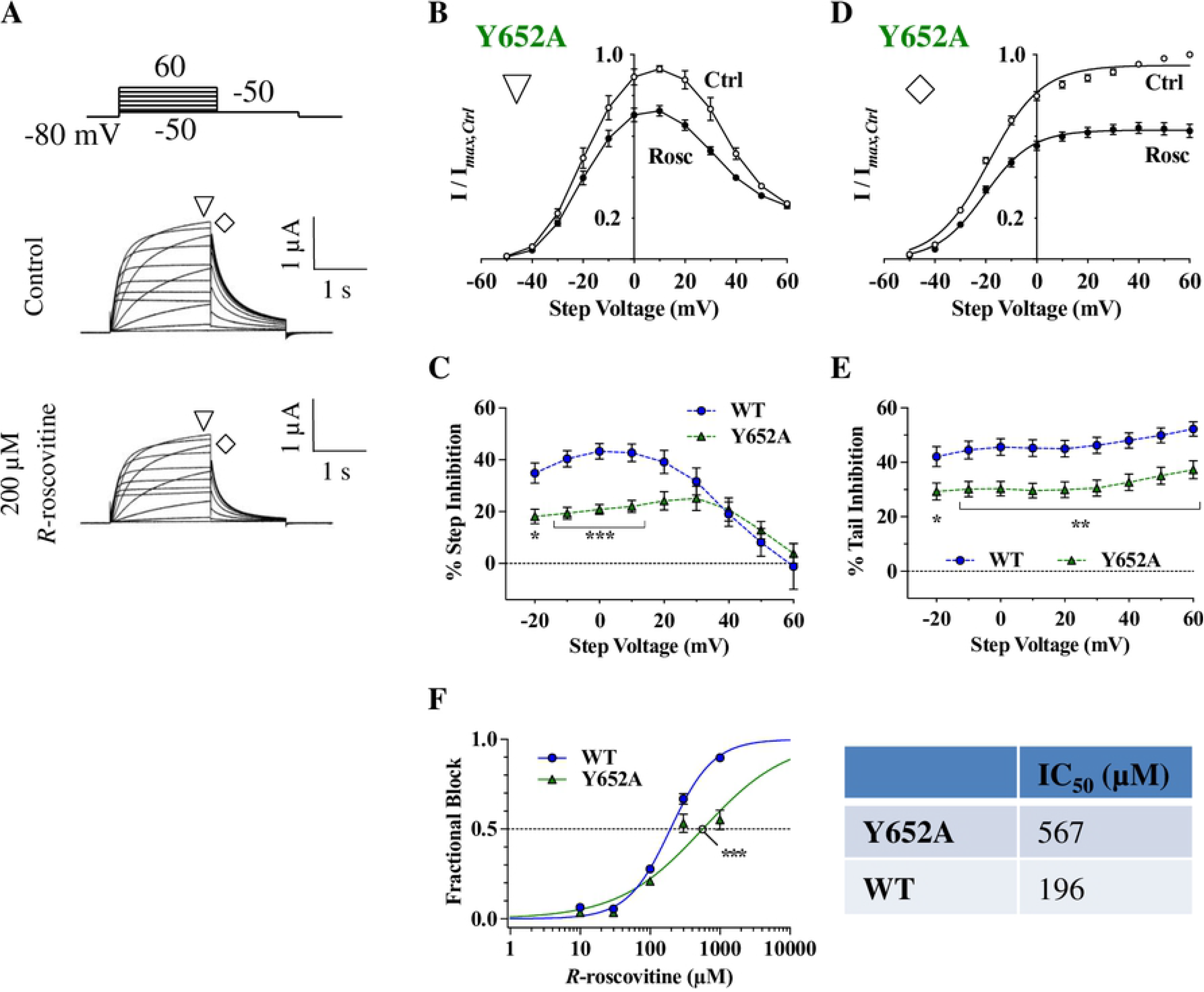
Y652A mutant hERG channels are resistant to inhibition by *R*-roscovitine. A) Voltage protocol and representative current traces of a cell before and during 200 μM *R*-roscovitine application. B) Y652A step I-V curves before and during 200 μM *R*-roscovitine inhibition (currents measured at ∇). The curves were n-shaped and displayed inhibition at intermediate voltages. C) Percent step inhibition of WT and Y652A. Levels of WT step inhibition were much stronger than that of Y652A between −20 mV and +10 mV (p < 0.05 for WT vs. Y652A, n_WT_= 14, n_Y652A_= 9, one-way ANOVA). D) Y652A tail I-V curves during the step depolarization protocol (currents measured at ♢). Boltzmann fits generated V_0.5_ (Ctrl: −16.5 ± 1.0 mV, Rosc: −19.6 ± 0.5 mV) and slopes (Ctrl: 12.5 ± 0.8, Rosc: 9.3 ± 0.3) of activation, with a small shift in V_0.5_ of activation having occurred (p = 0.0094, n = 11, paired *t*-test). E) Percent tail inhibition of WT and Y652A at their respective step voltages. Levels of Y652A tail current inhibition were significantly reduced compared to WT between −20 mV and +60 mV (p < 0.05 for WT vs. Y652A, n_WT_= 14, n_Y652A_= 9, one-way ANOVA). F) Concentration-response relationship for Y652A. When compared to WT (196 ± 12 μM), Y652A had an ~ 2.9-fold increase in IC_50_ (567 ± 122 μM) (p = 0.0005 for WT vs. Y652A, n_WT_= 11, n_Y652A_= 8, one-way ANOVA).

Analogous to WT tail I-V curves, Y652A tail I-V curves were shifted to more hyperpolarized voltages by *R*-roscovitine (V_0.5_ = −16.5 ± 1.0 mV vs. V_0.5,*Rosc*_ = −19.6 ± 0.5, p = 0.0094, n = 11, paired *t*-test; Fig 5D). Percent tail inhibition in Y652A was moderately reduced compared to WT between −20 mV and +60 mV, and showed the highest levels of inhibition at large depolarized potentials (p < 0.05 for WT vs. Y652A, n ≥ 9, one-way ANOVA; Fig 5E). Furthermore, the Y652A IC_50_ (567 ± 122 μM) was ~ 2.75-fold larger than WT (196 ± 12 μM; p = 0.0005 for WT vs. Y652A, n ≥ 8, one-way ANOVA; Fig 5F). The attenuated step and tail current inhibition of Y652A with 200 μM *R*-roscovitine, as well as the difference in IC_50_ values between Y652A and WT hERG, seemed to indicate that Y652 is involved in *R*-roscovitine binding. This was disimiliar to the results for S624A hERG (Fig 4), and made us question whether additional residues in the pore region were involved in *R*-roscovitine inhibition, and whether they would be more critical.

### The high K^+^ solution alters *R*-roscovitine inhibition of WT hERG

In addition to the residues S624 and Y652, we wanted to test whether other pore residues are more critical for inhibition: T623, a residue adjacent to the selectivity filter, and F656, an S6 residue, both of which are commonly associated with hERG block [32]. Mutating these two residues, however, alters permeation such that inward rectification is significant, with very little outward currents present. Thus, a high K^+^ solution (see methods), coupled with tail current measurements at −120 mV, were required to investigate the T623A and F656A hERG mutants [37,38].

With *R*-roscovitine binding to the hERG pore, it was likely that the large influx of K^+^ ions and the change in external solution would alter the WT hERG IC_50_. To determine the dose-response curves for *R*-roscovitine from WT tail currents, a pulsing protocol was applied, comprised of a 1 second depolarization to +40 mV and a 1 second repolarization to −120 mV; peak tail currents were measured in various *R*-roscovitine concentrations (Figs 6A, B). Fitting the fractional block of WT hERG tail current with a Hill equation resulted in an IC_50_ of 513 ± 43 μM (Fig 6C, n = 9), an ~ 2.6 fold larger value than that for the low K^+^ external solution (196 ± 12 μM). 500 μM *R*-roscovitine was, therefore, used with a step repolarization protocol to compare tail currents prior to and during drug application (Fig 6D). Measuring peak tail currents during repolarization shows WT hERG inhibition measured from the tail currents of the step repolarization protocol (see methods). Fitting tail I-V curves with quadratic equations shows that 500 μM *R*-roscovitine shifted the reversal potential from −11.1 ± 3.3 mV to −14.6 ± 2.4 mV (p = 0.0237 for Ctrl vs. Rosc, n = 10, paired *t*-test; Fig 6E; see methods), indicating a slight change in hERG permeation properties with *R*-roscovitine block. Furthermore, *R*-roscovitine inhibition levels increased as current direction changed from outward to inward: for example, WT hERG inhibition at both −120 mV (45.2 ± 2.3 %) and −80 mV (45.9 ± 2.5 %) was significantly larger than at +20 mV (27.0 ± 4.3 %; p < 0.0005 for either comparison; Fig 6F). The shift in *E*_rev_, along with the increased inhibition at hyperpolarized voltages, supported the idea that *R*-roscovitine binds to the hERG pore.

**Fig 6.**
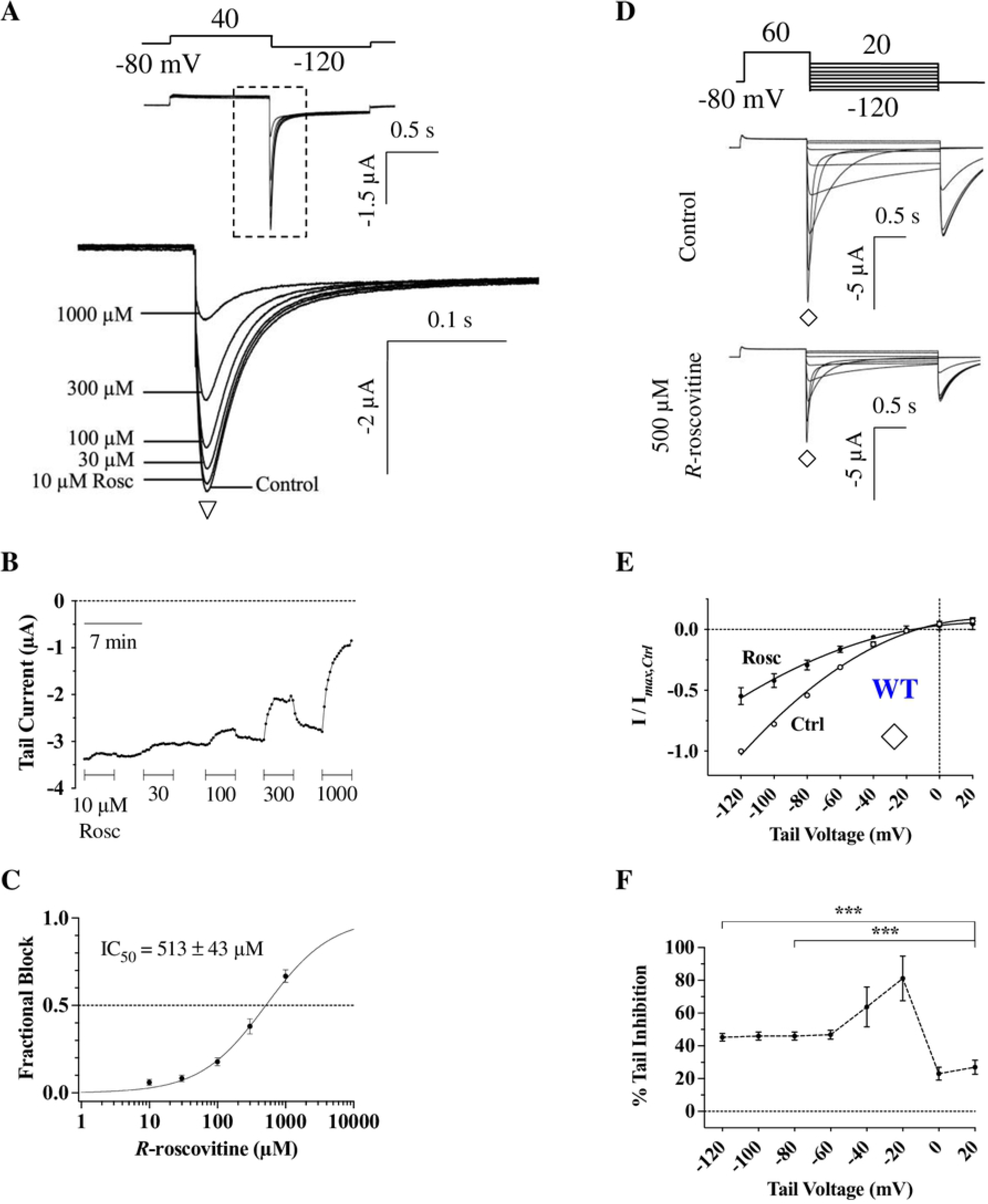
Inward WT hERG currents are inhibited by *R*-roscovitine. A) Pulse protocol with representative traces from a WT cell prior to (Control) and during various *R*-roscovitine contrations. Currents are superimposed in the diagram and peaks are magnified from the dashed box region. After a 1-second step to +40 mV from −80 mV, tail currents (∇) were elicited by a 1-second repolarization to −120 mV. B) Time course of inhibition from a representative cell. Peak tail currents were measured at ∇ and plotted over a span of ~ 36 minutes, during which different *R*-roscovitine concentrations were applied and washed off. C) Dose-response relationship for WT hERG in high K+. Fractional block data were fitted with a Hill equation to obtain the IC_50_ value (513 ± 43 μM, n = 9). D) Voltage protocol and representative current traces of a cell before (Control) and during 500 μM *R*-roscovitine application. E) WT tail I-V curves during the step repolarization protocol (currents measured at ♢). Smooth curves are quadratic fits, which were used to calculate reversal potential (*E*_rev_). 500 μM *R*-roscovitine shifted *E*_rev_ from −11.1 ± 3.3 mV to −14.6 ± 2.4 mV (p = 0.0237, n = 10, paired *t*-test). F) Percent tail inhibition of WT hERG at various repolarization voltages. Levels of inhibition were significantly stronger at low voltages than at high voltages (p = 0.0004 for +20 mV vs. −120 mV, p = 0.0002 for +20 mV vs. −80 mV, n = 10, one-way ANOVA).

### T623A and F656A hERG channels are insensitive to *R*-roscovitine

*R*-roscovitine exhibited a very weak inhibition of T623A hERG tail currents during a step repolarization protocol (Fig 7): between −60 mV and −120 mV, T623A inhibition was approximately half that of WT inhibition (p < 0.05 for WT vs. T623A, n = 10, one-way ANOVA). Consistent with reduced inhibition, plotting T623A tail currents against tail voltages resulted in quadratic nonlinear fits that showed no shifts in *E*_rev_ in the presence of *R*-roscovitine (Ctrl = −39.7 ± 2.0 mV, Rosc = −40.1 ± 2.1 mV; p = 0.5995 for Ctrl vs. Rosc, n = 10, paired *t*-test; Fig 7B). Furthermore, the attenuated potency of *R*-roscovitine with T623A hERG was supported by IC_50_ value comparisons: T623A IC_50_ was ~ 5.5-fold higher that that for WT (2786 ± 488 μM versus 513 ± 43 μM for WT, p = 0.0092, n ≥ 7, Kruskal-Wallis test; Fig 7D). The lack of an *E*_rev_ shift, along with the weakened inhibition and larger IC_50_ value compared to WT hERG, demonstrated the importance of T623 for *R*-roscovitine binding within the hERG pore.

**Fig 7.**
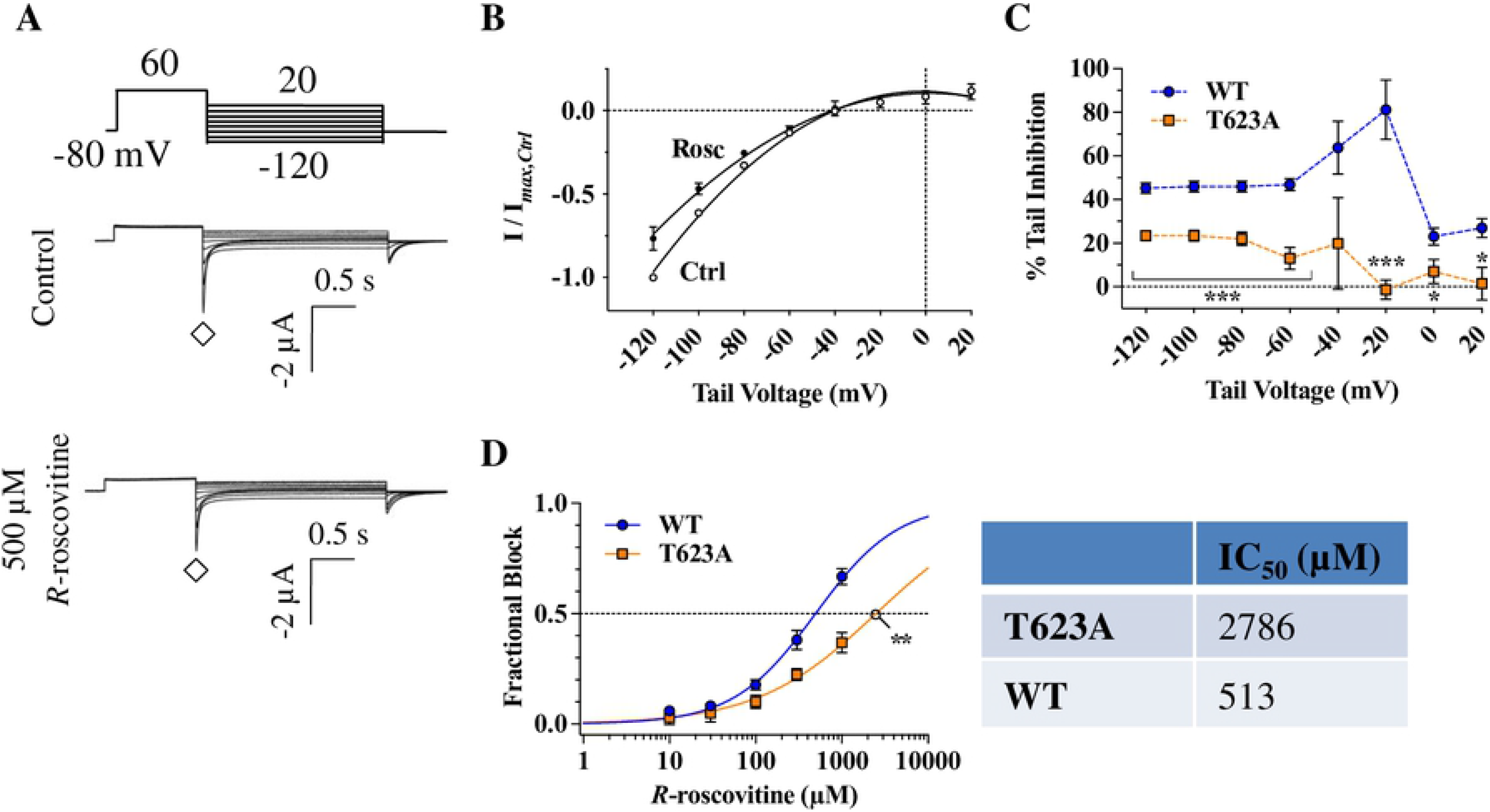
T623A hERG reduces *R*-roscovitine interaction within the pore. A) Voltage protocol and representative current traces of a cell before (Control) and during 500 μM *R*-roscovitine application. B) T623A tail I-V curves during the step repolarization protocol (currents measured at ♢). Quadratic fits generated *E*_rev_values (Ctrl = −39.7 ± 2.0 mV, Rosc = −40.1 ± 2.1 mV) that did not show a significant shift occurring with 500 μM *R*-roscovitine (p = 0.5995 for Ctrl vs. Rosc, n = 10, paired *t*-test). C) Percent tail inhibition of T623A hERG was significantly weaker over a range of voltages than WT inhibition (p < 0.05 for WT vs. T623A between +20 mV and −120 mV [excluding −40 mV, which is close to the reversal potential], n_WT_= 10, n_T623A_ = 10, one-way ANOVA). D) Compared to the WT IC_50_ (513 ± 43 μM), T623A IC_50_ was ~ 5.5-fold larger (p = 0.0092 for WT vs. T623A, n_WT_= 9, n_T623A_ = 7, Kruskal-Wallis test).

The F656A S6 mutant exhibited the largest levels of resistance to *R*-roscovitine. Tail currents of F656A hERG, in the presence of 500 μM *R*-roscovitine during a step repolarization protocol, were minimally impacted (Fig 8A). This initial observation extended to most tail voltages during repolarization (p > 0.05 for Ctrl vs. Rosc between +20 mV and −80 mV, n = 9, paired *t*-test; Fig 8B). At −120 mV, *R*-roscovitine inhibition of F656A hERG was only 12.3 ± 2.5 %, compared to the 45.2 ± 2.3 % inhibition of WT hERG (p < 0.0001 for WT vs. F656A, n ≥ 9, one-way ANOVA; Fig 8C). Surprisingly, there was a small but statistically significant shift in F656A *E*_rev_ (−45.1 ± 1.1 mV) by 500 μM *R*-roscovitine (−42.1 ± 1.7 mV; p = 0.02 for Ctrl vs. Rosc, n = 9, paired *t*-test). The IC_50_ for F656A was a ~ 42-fold increased from WT (21526 ± 10632 μM versus 513 ± 43 μM for WT, p = 0.0004, n ≥ 5, Kruskal-Wallis test; Fig 8D). These results, as well as the results for T623A hERG (Fig 7), indicated drastic modifications to the *R*-roscovitine binding site with mutations to either one of these hERG pore residues.

**Fig 8.**
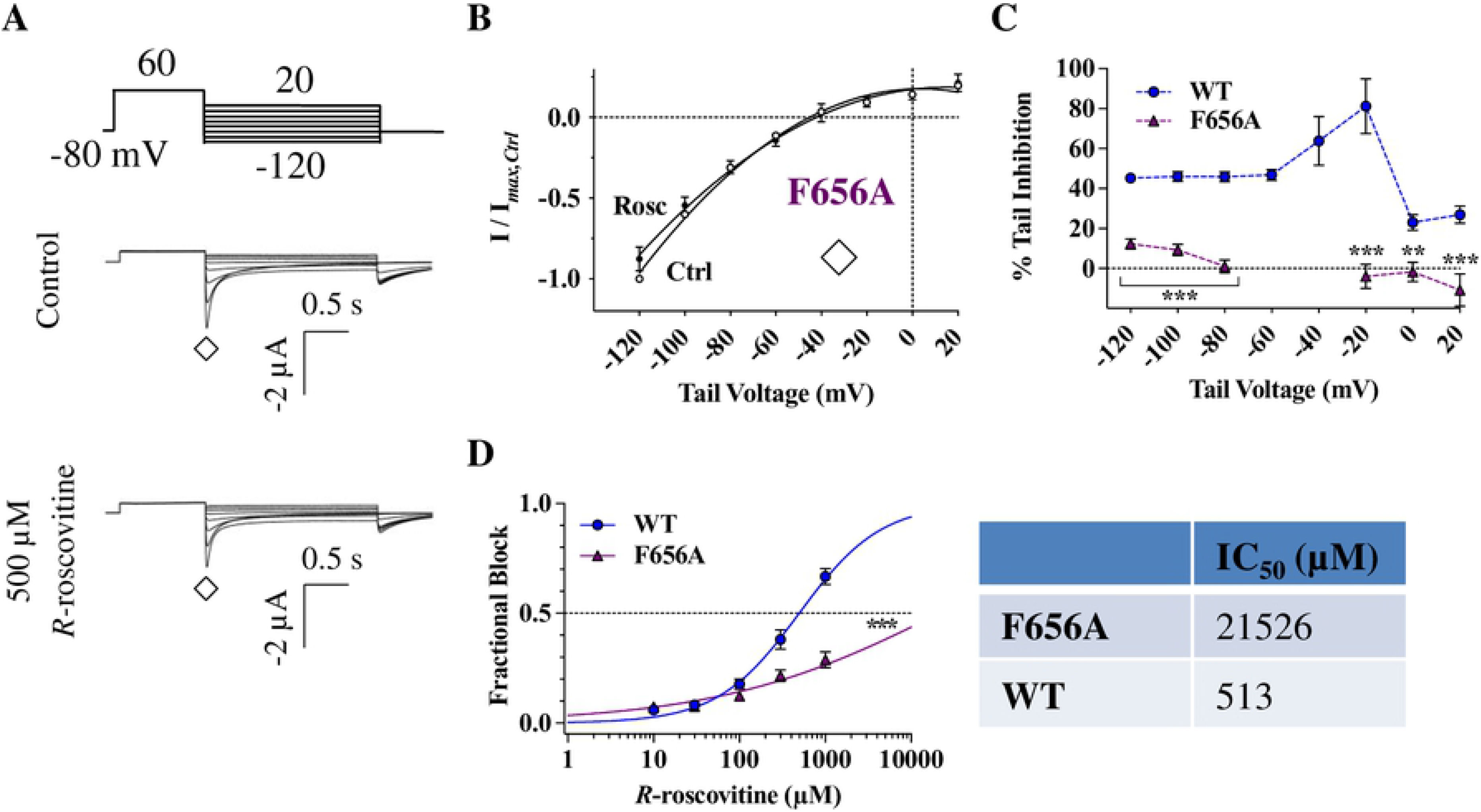
*R*-roscovitine inhibition is almost completely abolished in F656A hERG. A) Voltage protocol and representative current traces of a cell before (Control) and during 500 μM *R*-roscovitine application. B) F656A tail I-V curves during the step repolarization protocol (currents measured at ♢). Quadratic fits generated to obtain *E*_rev_ values (Ctrl = −45.1 ± 1.1 mV, Rosc = −42.1 ± 1.7 mV) did in fact show a shift with drug block (p = 0.02, n = 9, paired *t*-test). C) F656A tail current inhibition was almost non-existant for most repolarized voltages, which was a substantial difference from WT inhibition (p < 0.01 for WT vs. F656A between +20 mV and −120 mV [excluding −40 and −60 mV, which are both close to the reversal potential], n_WT_= 10, n_F656A_ = 9, one-way ANOVA). D) Concentration-response relationship for F656A hERG IC_50_ (21526 μM) shows a ~ 42-fold increase from WT IC_50_ (513 μM; p = 0.0004 for WT vs. F656A, n_WT_= 9, n_F656A_ = 5, Kruskal-Wallis test).

## Discussion

In this study we investigated the molecular determinants of *R*-roscovitine inhibition in the hERG pore, which has not been previously explored. *R*-roscovitine was expected to have a unique mechanism of inhibition when compared to other hERG inhibitors because of its preferential binding to open hERG channels and the properties outlined previously [25]. Our data suggests this indeed is the case since F656 and T623 are critical components of *R*-roscovitine inhibition, while S624 seems dispensable and Y652 contributes a relatively weak interaction. This is unlike the vast majority of other drugs for which the importance of these residues had been investigated. For example, MK-499 mediated hERG inhibition is weakened by a mutation to either S624A or Y652A [39], and ranolazine is heavily relient on both Y652 and F656 for inhibition [37]. A similar weakening of block with either S624, Y652, or F656 mutations were observed for many drugs, such as nifekalant [40], terfenadine, cisapride, [41], clofilium, ibutilide [42], bupivacaine [43], and propafenone [44], to name a few.

### The role of specific pore residues in hERG-*R*-roscovitine binding Y652A and F656A

The first S6 mutant to be characterized with *R*-roscovitine was Y652A hERG. 200 μM *R*-roscovitine inhibited both step and tail currents (Figs. 5A, 5B), as well as caused a negative shift in activation curves (Fig 5D). This shift in the presence of *R*-roscovitine, which is similar for both WT and Y652A, implicates *R*-roscovitine in open state stabilization/binding [45]. However, Y652A current inhibition levels were distinctly smaller than the inhibition of WT hERG (Figs. 5C, 5E), which corroborated other studies that identified Y652 as a critical residue for binding hERG inhibitors [26,36,39]. Nevertheless, the ~2.75-fold increase in IC_50_ value (Y652A IC_50_ = 567 ± 122 μM, WT IC_50_ = 196 ± 12 μM) meant that there is a weak but statistically significant leftover inhibition of Y652A tail current (Fig 5E), suggesting that *R*-roscovitine may still bind to the pore, but perhaps, in a different orientation, and warranted further investigation.

The other S6 mutant, F656A hERG, displayed a near complete insensitivity to *R*-roscovitine; F656A tail currents were almost indistinguishable before and during 500 μM *R*-roscovitine application (Figs 8A, 8B). The significant loss of inhibition, when compared to WT (Fig 8C), identified F656 as a critical residue for *R*-roscovitine binding. The larger value for F656A IC_50_ versus WT further confirmed the critical role of F656. However, in spite of the weakened affinity, there was a positvie shift in F656A *E*_rev_ during *R*-roscovitine application, suggesting that permeation is altered in the presence of *R*-roscovitine. This is in contrast to the lack of a *E*_rev_ shift with T623A hERG, a mutant that exhibited a very weak, but nonetheless apparent, current inhibition. The reasons for this remain unclear.

When comparing the two S6 mutant hERG channels (Y652A and F656A), we observed that the F656A mutation had a much more dramatic effect to abolish inhibition, as compared to the milder effect of the Y652A mutation. This was surprising because these residues have been previously shown to work in conjunction with one another to bind hERG inhibitors [37]. The divergence in the levels of Y652A and F656A inhibition could be due to the difference in binding interactions with *R*-roscovitine, with F656 primarily supplying a strong *π*-*π*/cation-*π* interaction [28,39]. Future *in silico* studies will investigate this. A previous study on mexiletine, a type Ib antiarrhythmic agent, had also found a stronger effect of the F656A compared to the Y652A hERG mutation to reduce inhibition (~28-fold versus ~12 fold reduction in IC50 in relation to WT, respectively), but it was proposed that both residues provide *π*-*π*/cation-*π* interactions [46]. In spite of the common molecular determinants, the two drugs appear to have different mechanisms of inhibition, with hERG activation and deactivation rates being affected by mexiletine inhibition but not by *R*-roscovitine inhibition [25]. It would be intriguing to further characterize interactions between mexiletine and residues in/adjacent to the selectivity filter (i.e. T623 and S624) for an even more extensive pharmacophoric analysis, which may shed light on the underlying preference of *R*-roscovitine for open state binding.

### T623A and S624A

Residues T623 and S624 are positioned deep within the hERG pore, specifically at the base of the pore helix [28]. With both side chains containing hydroxyl groups, hydrogen-bonding is frequently implicated in their interactions with drugs [39,42,47]. Depending on the orientation and structure of drugs that bind to the pore, mutating these pore residues can impact intermolecular interactions. This was evident with the T623A hERG mutant, which exhibited an overall weak tail current inhibition by *R*-roscovitine (Fig 7A). Upon analysis the shift in *E*_rev_ in the presence of *R*-roscovitine, it was apparent that the drug was not altering permeation (Fig 7B). Furthermore, the larger IC_50_ value for T623A hERG versus WT hERG suggested that the drug efficacy was reduced by the mutation (Fig 7D). Collectively, our data suggest the drug was no longer able to bind in its original pore location. This level of inhibition loss could represent *R*-roscovitine’s requirement of hydrogen-bonding to the hydroxyl group of T623 for maintaining normal potency.

Surprisingly, step and tail current inhibition were not impacted by mutation to the S624 residue (Figs 4B, 4D). In fact, inhibition levels increased in the S624A hERG mutant when compared to WT inhibition at the same voltages (Figs 4C, 4E), and no general increase in IC_50_ was apparent for S624A (Fig 4F). Collectively, these results suggest that S624 in WT hERG is an unlikely binding determinant. S624 has been previously implicated in the dissociation of hERG blockers: recovery from propafenone-mediated inhibition of hERG was significantly reduced in the S624A mutant [44] and, in contrast, S624A sped recovery from clofilium-mediated inhibition [42]. S624A hERG has also exhibited enhancement of inhibition with certain drug analogues [48]. Either way, this residue may be associated with *R*-roscovitine unbinding, which likely explains the increased inhibition in the S624A mutant.

In summary, our experimental results suggest that *R*-roscovitine has a unique binding orientation within the open WT hERG channel, relying on S6 residues (Y652, F656) and T623 but not S624 (a selectivity filter residue). This is, indeed, unlike many other hERG inhibitors mentioned above and likely contributes to its unique pharmacological mechanism of action.

### Physiological Relevance

Known hERG inhibitors linked to drug-induced LQTS, such as terfenadine, cisapride, astemizole, and sertindole, have been withdrawn from the market due to their inhibition of hERG channels [49]. The initial intrigue of *R*-roscovitine for this study was that a clinical trial using a range of *R*-roscovitine concentrations did not find QT prolongation or other cardiac side effects [20], such as those commonly seen with other hERG inhibitors. Thus, studies on *R*-roscovitine-hERG interactions may shed light on desirable drug features that could be used to identify potentially life-saving drugs that were never tested clinically due to hERG block.

One factor that likely contributes to the lack of arrhythmogenic side effects with *R*-roscovitine is its potency and, more importantly, it pleiotropic effects. *R*-roscovitine has a low affinity for hERG that may limit its adverse effects, even though the values used in this study and in other research are close to clinically relevant doses (10-30 μM) [50]. (*Xenopus* oocytes require higher doses than other cells because of their larger size and richer lipid component [51]). The relatively low affinity of *R*-roscovitine for hERG likely stems from its selective binding to the open state [25], as high affinity hERG inhibitors tend to bind to the inactivated hERG state [52]. Furthermore, this drug is unique when compared to class III antiarrhythmic drugs and other hERG inhibitors that, unlike *R*-roscovitine, exhibit: 1) binding to selectivity filter residues [18,39,53], 2) have relatively slow kinetics of recovery from inhibition, 3) are trapped in the hERG pore, and 4) exhibit use-dependence [25,54–56]. Our results provide a molecular basis for the observed unique *R*-roscovitine pharmacological properties, where the drug binds to the S6 residues (Y652 and F656) and a residue in close proximity to the selectivity filter (T623) but not S624, a selectivity filter residue.

Another key factor that may explain the lack of arrhythmogenic side effects could be the ‘multiple ion channel block effect’ [57], since *R*-roscovitine also inhibits L-type calcium channels [22]. Indeed, certain drugs bind to more than one voltage-gated ion channel, causing mutually-opposing actions on cardiac action potentials [58]. As a result, the re-classification of *R*-roscovitine as a Class VI antiarrhythmic agent was recently proposed due to its selective non-peak block of the late I_Ca,L_current, which likely averts cardiac arrhythmias [59,60]. Such compensatory mechanisms are now sought after, with the CiPA guidelines recommending that all drugs be screened not only against hERG but also for L-type calcium channel and sodium channel block [19]. Thus, revisiting drugs that were deemed clinically unsafe due to hERG block, but exhibit compensatory drug actions, may yield new pharmaceutical therapeutics and will support the CiPA initiative on proarrhythmia risk assessments.

## Acknowledgements

We thank Dr. Jian Yang (Columbia University) and Yong Yu (St. John’s University) for providing oocytes and technical expertise (J.Y) and for assistance and comments with the manuscript (Y.Y). For funding sources see financial disclosures section.

## Disclaimer

The content is solely the responsibility of the authors and does not necessarily represent the official views of the National Institutes of Health.

